# *Elizabethkingia anophelis* response to iron stress: physiologic, genomic, and transcriptomic analyses

**DOI:** 10.1101/679894

**Authors:** Shicheng Chen, Benjamin K. Johnson, Ting Yu, Brooke N. Nelson, Edward D. Walker

## Abstract

*Elizabethkingia anophelis* bacteria encounter fluxes of iron in the midgut of mosquitoes, where they live as symbionts. They also establish bacteremia with severe clinical manifestations in humans, and live in water service lines in hospitals. In this study, we investigated the global gene expression responses of *E. anophelis* to iron fluxes in the midgut of female *Anopheles stephensi* mosquitoes fed sucrose or blood, and in iron-poor or iron-rich culture conditions. Of 3,686 transcripts revealed by RNAseq technology, 218 were upregulated while 112 were down-regulated under iron-poor conditions. Most of these differentially expressed genes (DEGs) were enriched in functional groups assigned within “biological process,” “cell component” and “molecular function” categories. *E. anophelis* possessed 4 iron/heme acquisition systems. Hemolysin gene expression was significantly repressed when cells were grown under iron-rich or high temperature (37°C) conditions. Furthermore, hemolysin gene expression was down-regulated after a blood meal, indicating that *E. anophelis* cells responded to excess iron and its associated physiological stress by limiting iron loading. By contrast, genes encoding respiratory chain proteins were up-regulated under iron-rich conditions, allowing these iron-containing proteins to chelate intracellular free iron. *In vivo* studies showed that growth of *E. anophelis* cells increased 3-fold in blood-fed mosquitoes over those in sucrose-fed ones. Deletion of aerobactin synthesis genes led to impaired cell growth in both iron-rich and iron-poor media. Mutants showed more susceptibility to H_2_O_2_ toxicity and less biofilm formation than did wild-type cells. Mosquitoes with *E. anophelis* experimentally colonized in their guts produced more eggs than did those treated with erythromycin or left unmanipulated, as controls. Results reveal that *E. anophelis* bacteria respond to varying iron concentration in the mosquito gut, harvest iron while fending off iron-associated stress, contribute to lysis of red blood cells, and positively influence mosquito host fecundity.

## Introduction

*Elizabethkingia anophelis* is an aerobic, non-fermenting, Gram-negative rod (1). It is ubiquitously distributed in diverse natural environments including water, soil, sediment, plants, and animal digestive tracts (2, 3). *E. anophelis* associates symbiotically with the gut lumen environment of *Anopheles* (1) and *Aedes* mosquitoes, whether in wild-caught individuals or those from insectary colonies (4, 5). *E. anophelis* enriched in the larval *Anopheles* gut during filter feeding from the surrounding water medium, transmitted transtadially from larval to adult gut lumen during metamorphosis, and transmitted vertically to the next generation through an uncharacterized mechanism (3). *E. anophelis* infection resulted in high mortality in adult *Anopheles* when injected through the cuticle into the mosquito hemocoel, exhibiting pathogenesis, but stabilized as nonpathogenic symbionts when fed to the mosquitoes and confined to gut (6). These findings indicate that *E. anophelis* can be opportunistically pathogenic in mosquitoes but that the gut provides a barrier against systemic infection (6). Hospital-acquired infections of *E. anophelis* occur in sick or immunocompromised individuals, sometimes leading to death (2, 7, 8). *E. anophelis* was the cause of a recent outbreak of nosocomial illness in the Upper Midwest region of the United States (Wisconsin, Illinois and Michigan) (2, 9, 10). Other outbreaks have occurred in Africa, Singapore, Taiwan and Hong Kong (8, 11, 12). A survey of five hospitals in Hong Kong showed that bacteremia in patients due to *E. anophelis* was often associated with severe clinical outcomes including mortality (11). Bacteria disseminated from hospital water service lines, especially sink faucets, during handwashing to the hands of healthcare workers, who subsequently exposed patients when providing care (13). The association between bacterial infection in mosquitoes and infection in human beings is not established (14).

Regular but episodic influxes of blood enter the female mosquito midgut, greatly but temporarily altering gut physiology and environmental conditions, including an increase in proteolytic enzyme activity associated with blood meal digestion, formation of a chitinous peritrophic matrix around the blood meal, a sudden and large increase in temperature, and large flux of iron from heme and ferric-transferrin sources (15, 16). Mosquitoes also take sugar meals, whose first destination is the foregut “crop” from where sugar moves to the midgut for digestion and assimilation (17). Iron is practically nil in the midgut after a sugar meal and between blood meals, but spikes to ca. 600 ng iron per ul after a blood meal (18). The community structure of the microbiome in the mosquito gut varies temporally with episodes of blood and sugar feeding (3, 19). Our specific interest in *E. anophelis* focuses on understanding the survival mechanisms and physiological adaptations as it establishes and maintains infection in the mosquito gut, where environmental conditions such as iron flux are highly variable.

Iron is an essential cofactor in many enzymes that are involved in maintaining cell homeostasis and functions (20). Thus, bioavailability of iron greatly influences bacterial metabolism, growth and transcription (21-23). In the mosquito midgut, the microbiota meet two very different iron concentrations. Very limited iron (non-heme form, Fe^3+^ or Fe^2+^) will be available in the female midgut when only nectar is imbibed (18). Microorganisms might employ various mechanisms to scavenge iron under those conditions (23). One of the most efficient strategies to sequester iron is to secrete iron chelator siderophores (24). Iron-bound siderophores are transported into periplasm through specific siderophore receptors (TonB-dependent iron transports) or other transport systems (25). Heme/iron increases suddenly when a mosquito ingests and digests erythrocytes, increasing oxidative stress in the gut lumen (26, 27). Bacteria secrete hemophores to capture hemin from hemoproteins (released from hemoglobin when erythrocytes are disrupted) and deliver it to bacterial periplasm (28).

The ability to scavenge iron, manage iron-induced stress, and minimize damage from reactive oxygen radicals generated by iron could influence bacterial colonization and survivorship in the mosquito gut (24, 25, 29). In this study, we hypothesize that gene expression revealing these processes in *E. anophelis* will be significantly influenced by iron availability in the mosquito midgut, depending on the two normal types of meals (blood vs sugar). To explore this hypothesis, we analyzed global transcriptomic changes in *E. anophelis* under iron-replete (or iron-rich) and iron-depleted (or iron-poor) conditions. We developed a genetic manipulation system to knock out siderophore synthesis genes whose expression was significantly regulated by iron. Furthermore, we labeled wild-type and mutant bacteria with sensitive luciferase-based reporters for purposes of quantifying their growth and gene expression in mosquitoes. Hemolysin genes responding to iron availability *in vitro* and blood meals *in vivo* were investigated in detail. The goal was to use a combination of transcriptomic and genetic analyses of iron metabolism in *E. anophelis* to expand our understanding of bacterial survival mechanisms and physiological functions in the mosquito gut.

## Materials and Methods

### Bacterial strains, growth conditions, and molecular manipulations

Strains and molecular reagents used in this study are listed in Table 1. *E. coli* DH5α was used for cloning. *E. coli* S17 (*λ pir*) was used for conjugation. Luria-Bertani (LB) media were used for *E. coli* cultures. *E. anophelis* Ag1 was isolated from the mosquito *An. gambiae* (30) and was cultured in tryptic soy broth (TSB) or LB media (31). Liquid cultures were grown with shaking (*ca*. 200 rpm) at either 30°C (*E. anophelis* and *F. johnsoniae*) or 37°C (*E. coli*). For solid LB media, Bacto-Agar (Difco, Detroit, Michigan) was added to a final concentration of 20 g/L. Whenever necessary, erythromycin (100 µg/ml) (abbreviation, Em), kanamycin (50 µg/ml) (abbreviation, Km) or ampicillin (100 µg/ml) (abbreviation, Amp) was added to media to screen transconjugants or bacteria with plasmids, respectively.

**Table 1.**
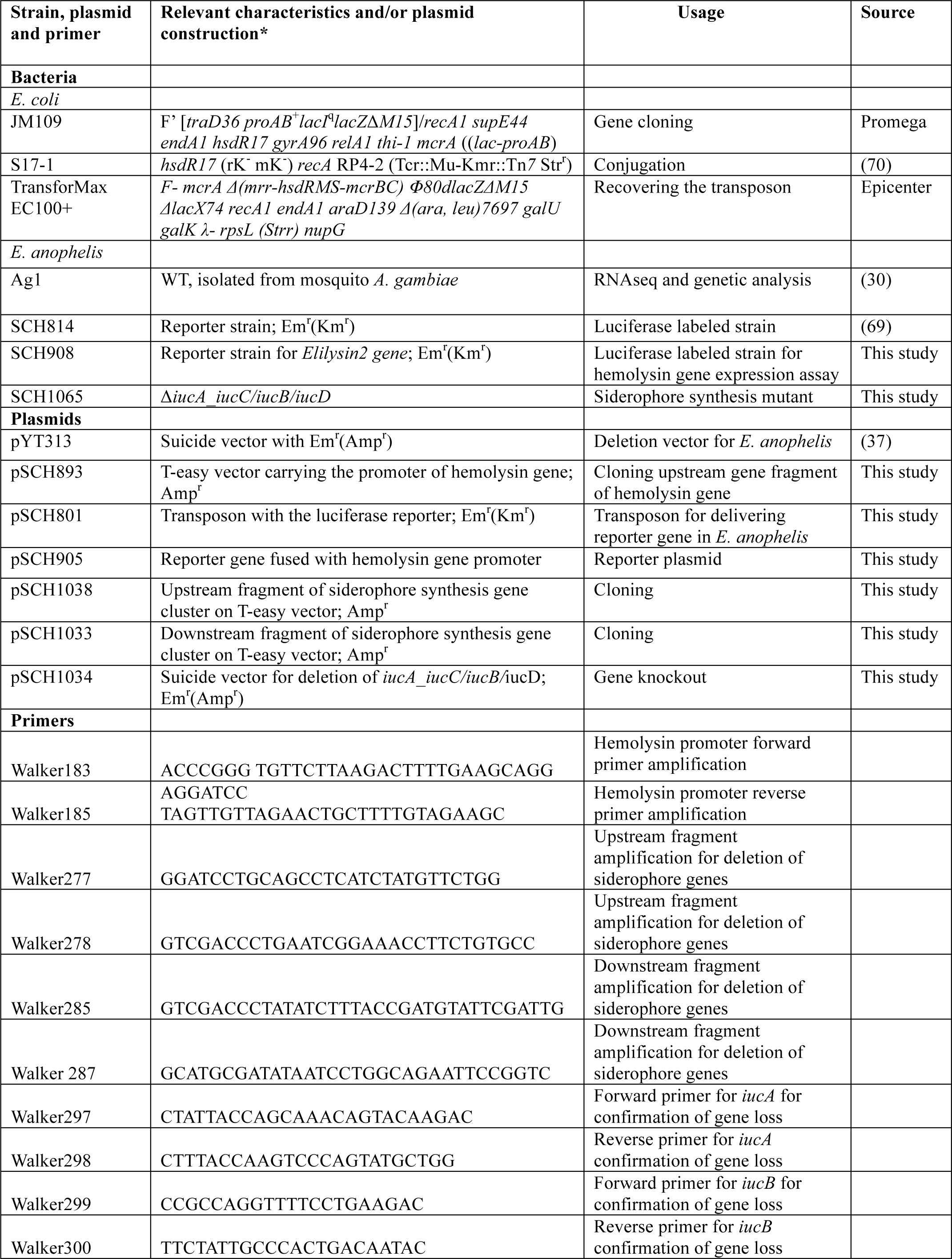
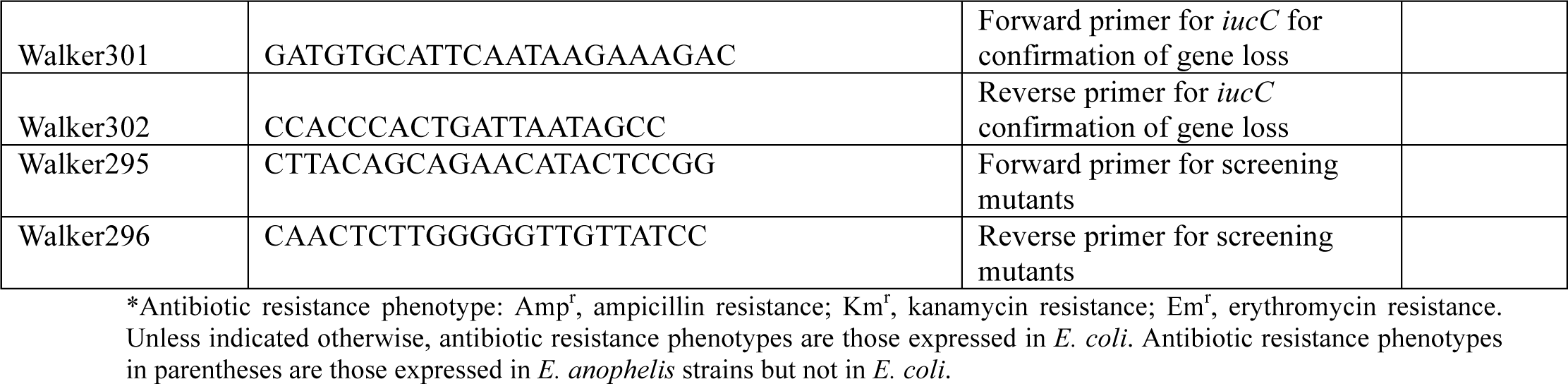
Strains, plasmids and primers used in this study.

Genomic DNA was prepared using a genomic DNA extraction kit (Promega, Madison, WI, USA), and plasmid DNA was purified with the QIAprep spin miniprep kit (QIAGEN, Germantown, MD, USA). PCR amplifications were done with the Failsafe PCR system (Epicenter Technology, Madison, WI, USA). Amplicons were separated in 0.7–1.0% (w/v) agarose gels, and DNA fragments were purified with the QIAquick gel extraction system (QIAGEN). Restriction and modification enzymes were purchased from Promega (Madison, WI, USA) or New England Biolabs (Beverly, MA, USA). Ligation mixtures were transformed into *E. coli* cells, and transformants were plated onto LB plates with appropriate antibiotic selection. Resistant colonies were isolated, and then screened for the acquisition of plasmids. All constructs were sequenced to verify the structure.

### RNA preparation and Illumina RNA-seq

*E. anophelis* cultures were grown overnight in LB and OD_600nm_ was adjusted to 0.1 by diluting the cells into 20 ml of LB supplemented with either 12 µM of FeCl_3_ or 200 µM of 2, 2-dipyridyl in 250 ml flasks. Cells were grown at 30°C with shaking (200 rpm) for 3 hours to a final OD of 0.2 or 0.5 for iron-depleted cultures or iron-replete ones, respectively. Cells grown in log-phase were re-suspended in RNAlater and frozen immediately after being harvested. RNA was isolated using Trizol reagents following the manufacturer’s manual. Residual DNA in the samples was removed using Dnase I. The integrity of the RNA was analyzed using an Agilent bioanalyzer (Agilent Technologies). The Ribo-Zero rRNA removal kit (Gram-negative bacteria, Epicentre) was used to remove the ribosomal RNA species (23S and 16S rRNA) from total RNA in samples. Library construction and sequencing were performed by Beijing Genomics Institute (BGI) using TruSeq RNA sample preparation v2 guide (Illumina). Three biological replicates of each treatment were used for RNA-Seq. The libraries were sequenced using the Illumina HiSeq 2000 platform with a paired-end protocol and read lengths of 100 bps.

### RNA-seq data analysis

Data from RNA-seq were checked for quality control (QC), pre-trimming, using Fast QC (https://www.bioinformatics.babraham.ac.uk/projects/fastqc/). Raw reads were subjected to trimming of low-quality bases and removal of adapter sequences using Trimmomatic (v0.32) with a 4-bp sliding window (32) when the read quality was below 15 (using the Phred64 quality scoring system) or read length was less than 50 bp. The trimming process improved the quality of the data as evidenced by comparing Fast QC reports pre- and post-trimming. Forward and reverse read pairs were aligned to the reference genome using Bowtie2 (v2.2.3) with the –S option to produce SAM files as output (http://bowtie-bio.sourceforge.net/bowtie2/index.shtml). SAM files were converted to BAM format, sorted by name using the –n option and converted back to SAM format using SAMTools (33). Aligned reads were then counted per gene feature in the *E. anophelis* Ag1 genome using the HTSeq Python library (34). Specifically, counting was performed using the htseq-count function within the HTSeq suite of tools using the “–r” name and “–s” no options. Differential gene expression was calculated by normalizing the data utilizing the trimmed mean of M-values normalization method and filtering out genes that had <10 counts per million (CPM) within the edgeR package (35). Statistical analysis was performed in RStudio (v 0.98.1102) (36) by the exact test with a negative binomial distribution for each set of conditions and testing for differential gene expression using edgeR (35). Differentially expressed genes were determined to be statistically significant based on an adjusted p < 0.05. Magnitude amplitude (MA) plots were generated by modifying a function within the edgeR package (35). Red dots indicate statistically significantly, differentially expressed genes (adjusted p < 0.05) and black dots are non-statistically significantly, differentially regulated genes. Blue lines indicate two-fold changes either up- or down-regulated.

### Reporter system for hemolysin gene expression in *E. anophelis*

*E. anophelis* possessed at least three genes encoding putative hemolysins (here named Elilysin1, EAAG1_11032; Elilysin2, EAAG1_11027; Elilysin3, EAAG1_18561) (see Results). The promoter of the *Elilysin2* gene (EAAG1_11027) was chosen for the following experiment because it carried a typical promoter motif conserved in Bacteroidetes (Figure S1). Identification of the transcriptional start site, promoter, and regulatory region prediction is described in Figure S1. A 757-bp fragment spanning the 5’-end of EAAG1_11027 and the 3’-end of a hypothetical protein was amplified with primers Walker183 and Walker185 using genomic DNA as template. The amplicon was cloned into the T-easy vector (pSCH893) (Table 1). The insert was released from pSCH893 by restriction enzymes SmaI and SacII and ligated into the same sites on pSCH801, creating pSCH905 (Table 1). pSCH905 was conjugatively transferred into *E. anophelis* and colonies with erythromycin resistance and luciferase production were selected, leading to the reporter strain pSCH908 (Table 1).

### Deletion of genes encoding the siderophore synthesis cluster

The siderophore synthesis gene cluster consisting of *iucA/iucC* (EAAG1_10367), *iucB* (EAAG1_10372) and *iucD* (EAAG1_10377) was targeted for deletion. To accomplish it, upstream (1963-bp) and downstream (1657-bp) gene fragments were amplified with primers Walker277/Walker278 and Walker285/Walker 287, respectively, by using *E. anophelis* genomic DNA as the template. Amplicons were gel purified and separately cloned into pGEM-T easy (pSCH1038 and pSCH1033). Upstream fragments were released by BamHI/SalI digestion, and downstream fragments were released by SalI/SphI from pGEM-T easy and then sequentially assembled at the same sites on the suicide plasmid pYT313 (pSCH1034).

Plasmid pSCH1034 was conjugatively transferred into *E. anophelis* through procedures described elsewhere (37). Merodiploids were selected on LB plates supplemented with Em. The Em-resistant merodiploids were resolved by plating single colonies onto LB agar medium containing 10% (wt/vol) sucrose and Em according to a previously described method (37). Putative (Δ*iucA_iucC/iucB/iucD*) clones were identified by screening with PCR with primers Walker295/Walker296 (Table 1) and checked for susceptibility to Em and sucrose; one confirmed (Δ*iucA_iucC/iucB/iucD*) clone (SCH1065) was chosen for further analysis.

### Siderophore activity determination by CAS liquid assay

The CAS solution was prepared by following procedures previously described (38). The CAS solution consisted of HDTMA (0.6 mM), FeCl_3_ (15 µm), CAS (0.15 mM) and piperazine (500 mM). The buffer system was PIPES (1mM, pH 5.6). Due to the lack of the commercially available aerobactin as a reference, we utilized the purified deferoxamine mesylate salt (Sigma-Aldrich, USA) as a standard to measure siderophore activity. CAS solution was mixed with the equivalent volume of the filtered supernatants from various cultures, incubated at 37°C for 3 h, and the OD_630nm_ determined on a microplate reader.

### Hydrogen peroxide susceptibility assays

*E. anophelis* cultures grown overnight (16 h) were diluted 10-fold in 20 ml of LB supplemented with either 40 µM FeCl_3_, 10 µM hemoglobin, or 1.28 mM 2, 2-dipyridyl in a 250 ml flask. After 3 h, the log-phase cells were harvested and washed with 1 X PBS. After OD_600nm_ was adjusted to 0.1, the washed *E. anophelis* cells were treated with 20 µM of H_2_O_2_ (final concentration) for 20 min. Then cells were extensively washed with 1 X PBS three times before plating on LB agar for viable counts.

### Biofilm formation and quantification

*E. anophelis* was cultured in TSB broth at 37°C with agitation. Cultures grown overnight were diluted in TSB broth supplemented with either 40 µM of FeCl_3_ or 1.28 mM of 2, 2-dipyridyl, respectively. 200 µl of the above bacterial suspension was inoculated into individual wells of the 96-well polystyrene microtiter plates and statistically cultured overnight. TSB broth without bacterial inoculation was used as the negative control. A modified biofilm assay was carried out according to published methods (39, 40). Planktonic cells were removed and the absorbed cells were air dried for 30 min at room temperature. 0.5% crystal violet was added into the wells, incubated for 15 min at room temperature, and rinsed thoroughly with distilled water. After air drying, crystal violet was solubilized in 200 µl of ethanol acetone (80:20, vol/vol) for 30 min, and the OD_570nm_ was measured by using a SpectraMax M5 microplate reader (Molecular Devices, Sunnyvale, CA). 12 replicates were used for each treatment.

### Nucleotide sequence accession numbers

The RNA-seq data has been submitted to the NCBI Gene Expression Omnibus (GEO) under accession number GSE132933, to the NCBI BioProject under accession number PRJNA549490 and to the NCBI The Sequence Read Archive (SRA) under the accession number SRP201789.

## Results

### General transcriptome features

The raw sequence output of the transcriptomes included 150 million reads in total. Reads were well matched (100%) to the published *E. anophelis* Ag1 genome (3). Out of the 3,686 transcribed genes detected in this study (Table S1), 330 displayed a significant change (more than 4-fold, adjusted *P* value < 0.01), which counted for 9% of total transcripts in *E. anophelis* (Table S2). Among the significantly regulated genes by iron, 218 transcripts displayed a significant decrease (Figure 1A and 1B, Table S3) while 112 showed a significant increase (Figure 1A and 1B, Table S4). The remaining genes (n = 3356, i.e., ∼91% of total genes in *E. anophelis* transcriptomes) were non-DEGs (Table S1). Multidimensional scaling analysis of the matrix of up- and down-regulated genes by experimental category of high- or low-iron treatment samples showed that the transcriptomes of the biological replicates in iron-rich samples grouped together while they were well separated from those grown under iron-poor conditions (Figure S2).

**Figure 1.**
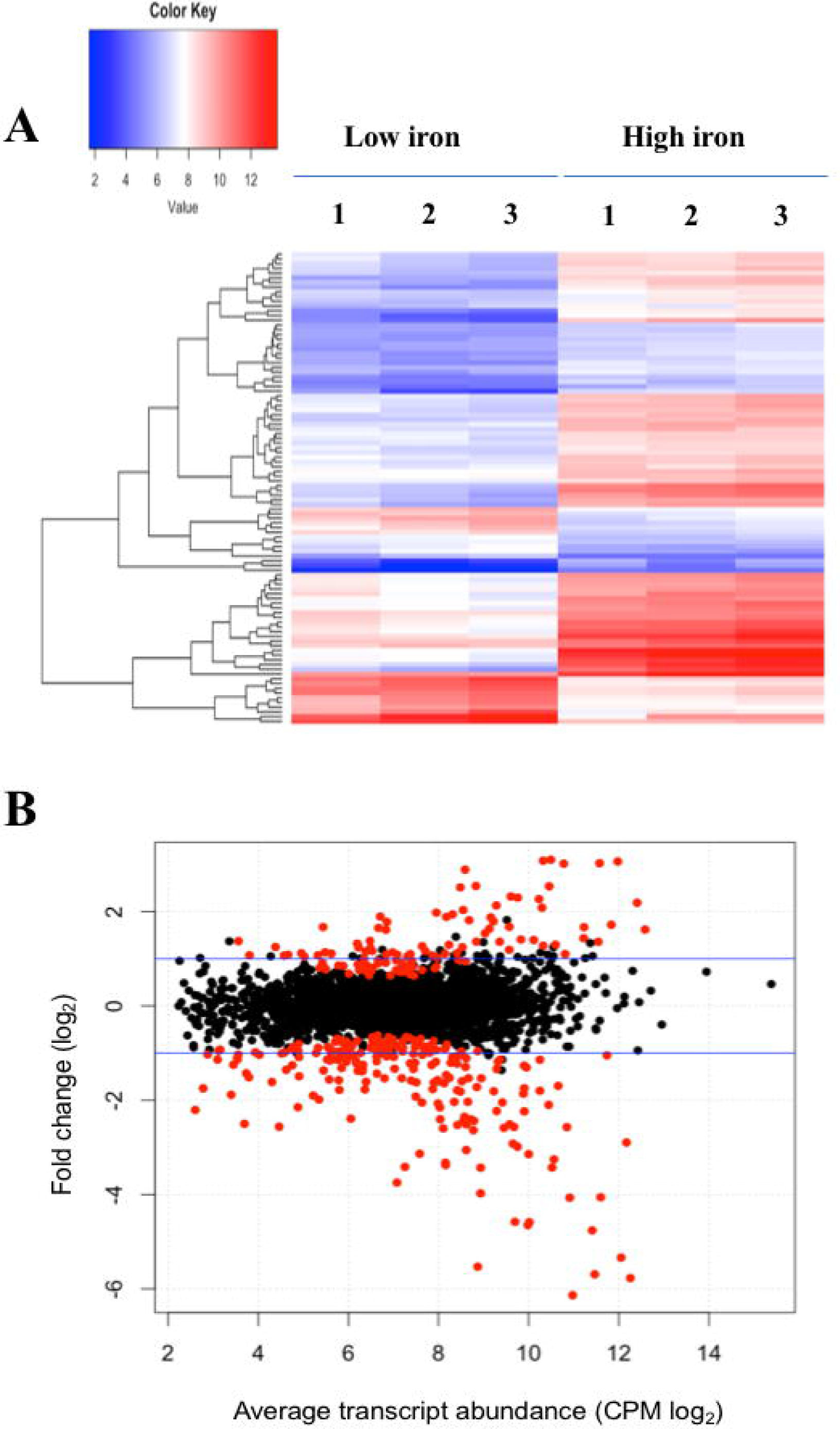
Comparison of differential gene expression between *E. anophelis* Ag1 cultures held in low- and high-iron culture conditions. (A) Heat maps of 100 genes with significant regulation by iron availability, with cluster analysis of gene relatedness. Left, high-iron condition; right, low-iron condition. (B) Magnitude amplitude plots generated by a modifying function within the edgeR package. Red dots indicate statistically significant genes (adjusted p < 0.05) and black dots are non-statistically significant differentially regulated genes. Blue lines indicate two-fold changes either up-regulated or down-regulated.

Enrichment analysis by Gene Ontology (GO) showed that the DEGs were assigned to at least 30 functional groups within 3 main GO categories including “biological process”, “cell component,” and “molecular function” (Figure 2). In the “cell component” category, genes encoding the respiratory chain complex, outer membrane, cell periphery and external encapsulating structure were notably up- or down-regulated by iron. Further, genes of the respiratory electron transport chain/complex were enriched in the “biological process” category. In the “molecular function” category, the functional groups were associated with oxidoreductase activity (acting on NAD(P)H), NADH dehydrogenases, and heme or quinone binding proteins (Figure 2). Enrichment analysis of KEGG pathways (Figure 3A) showed that the majority of up-regulated DEGs was assigned to “energy metabolism” (22) including oxidative phosphorylation (NADH-quinone oxidoreductase). Next to “energy metabolism”, the enriched DEGs were involved in “environmental information” (8) or “genetic information processing” (6). In contrast to up-regulated DEGs, down-regulated genes (Figure 3B) were predominantly enriched in the “genetic information” (25) and “environmental information” (21) processing categories, including translation and amino acid metabolism. Further, metabolic analyses by STRING (Figure 3C and 3D) indicated that many genes involved in oxidative phosphorylation (e.g. NADH:quinone oxidoreductases) were clustered together in high iron cells (Figure 3C). It is very likely that they interact with each other due to similar functions and cellular compartments. Many gene products involved in lipid, porphyrin and phenylalanine metabolism also formed clusters with possible interactions (Figure 3D). By contrast, amino acid synthesis (aromatic amino acid biosynthesis), iron uptake (siderophore, enterobactin and receptors), vitamin (B12) synthesis, ribosome proteins and thioredoxin were enriched and clustered together in iron-restricted cells (Figure 3D).

**Figure 2.**
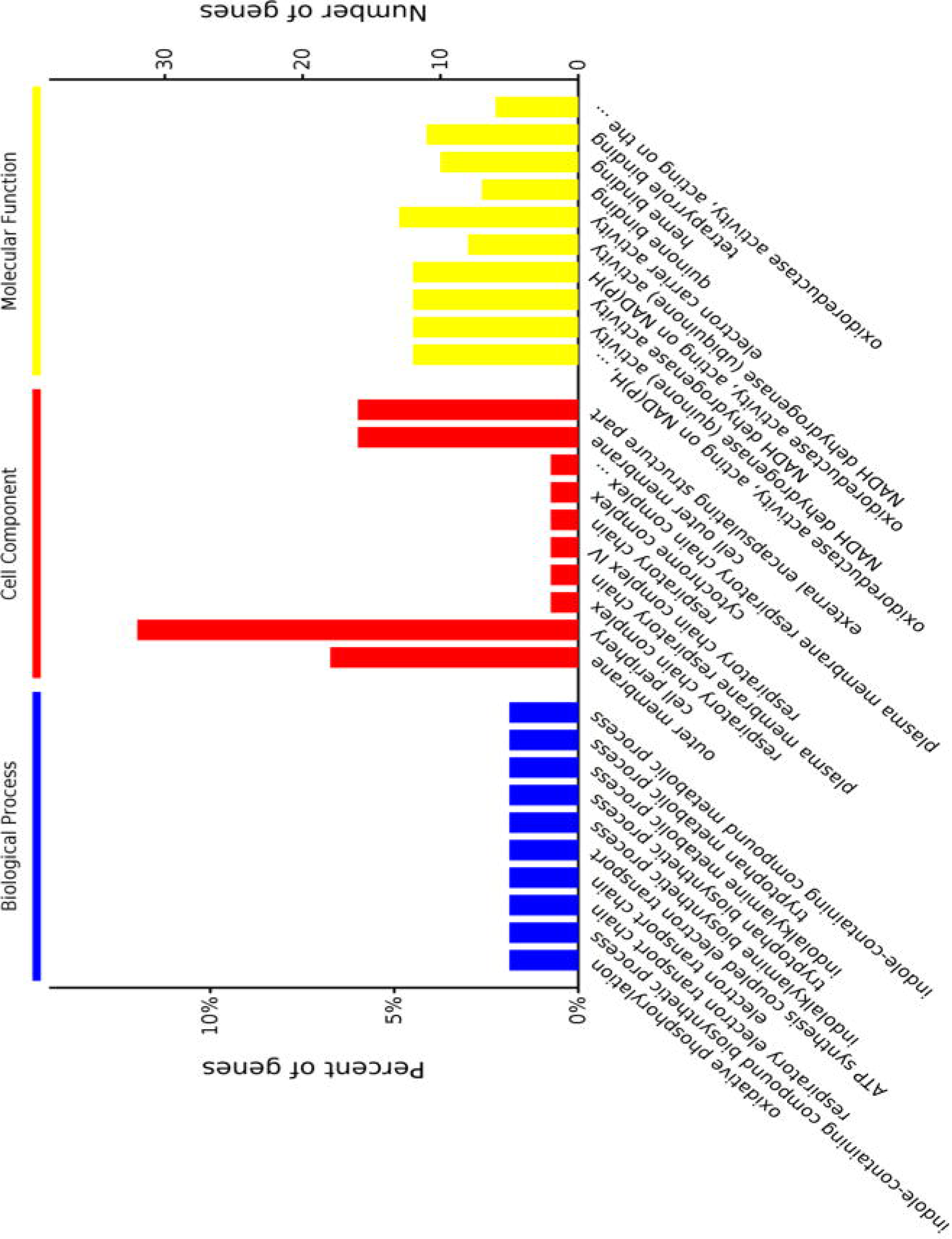
Distribution of the three main gene organization categories for differentially expressed genes in *E. anophelis* strain encountered in this study. The categories “biological process,” “cell compartment,” and “molecular function” are indicated with functional subcategories noted within each. The differentially expressed genes (adjusted *P* < 0.05) between high iron and low iron cells were identified by using the edgeR (v3.10.5).

**Figure 3.**
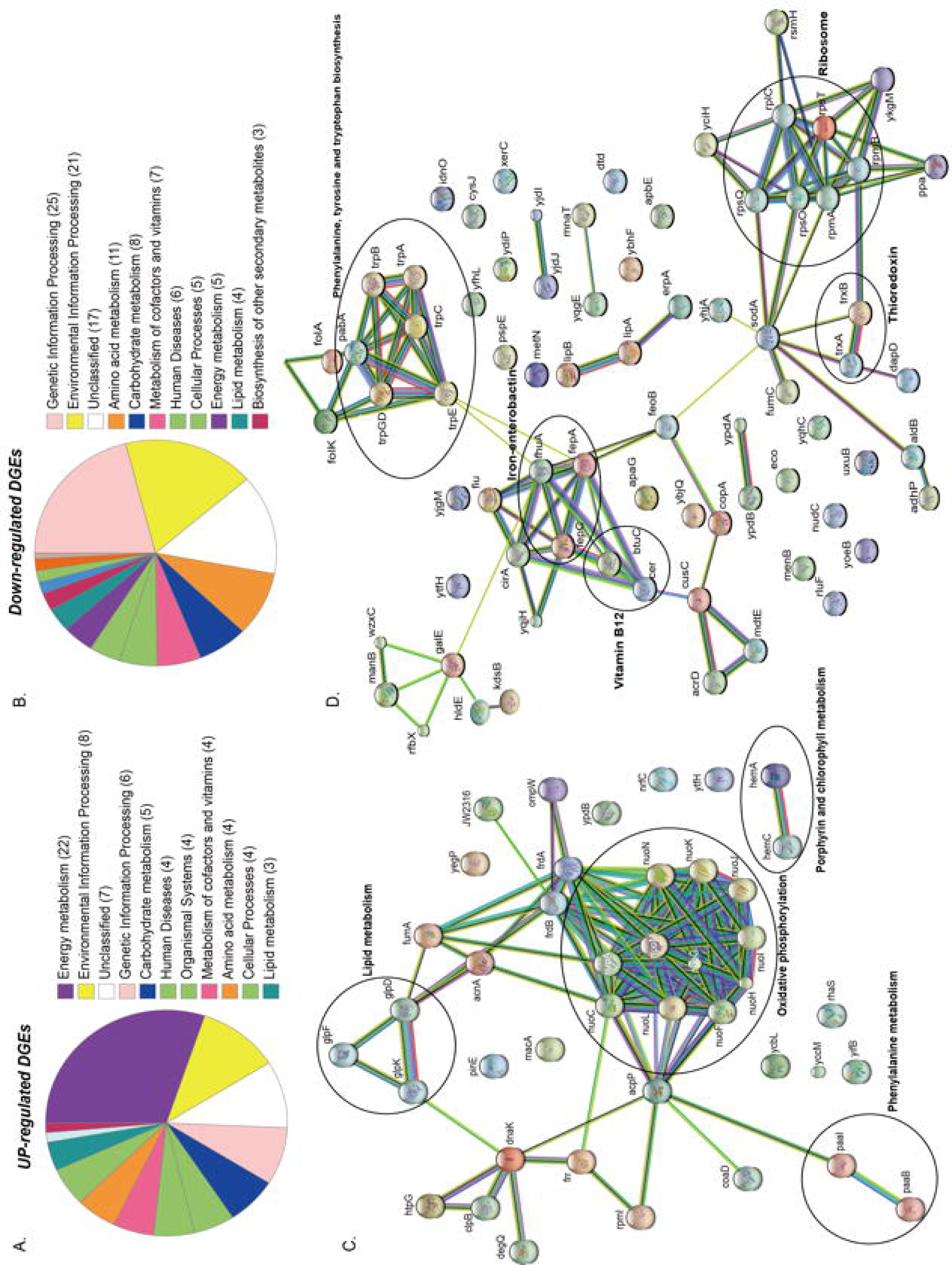
KEGG and STRING analysis of differentially regulated genes in *E. anophelis* Ag1. (A) KEGG pathway enrichment analysis of up-regulated DGEs. Most of up-regulated DGEs were enriched to Energy Metabolism. (B) KEGG pathway enrichment analysis of down-regulated DGEs. Most of up-regulated DGEs were enriched to Translation and Amino acid metabolism. (C) STRING analysis of up-regulated DGEs. Up-regulated DGEs were clustered to oxidative phosphorylation (circled). (D) STRING analysis of down-regulated DGEs. Down-regulated DEGs were clustered to ribosome and the biosynthesis of phenylalanine, tyrosine and tryptophan (circled).

### Respiratory chain complex in response to iron availability

Genes encoding the respiratory chain protein components were significantly up-regulated in iron-replete media (Table 2). Related to this, a gene cluster related to quinol:cytochrome c oxidoreductase (*bc1* complex or complex III) synthesis and assembly was newly discovered in *E. anophelis* (Figure 4A). The *bc1* cluster consists of cytochrome c2, quinol:cytochrome c oxidoreductase iron-sulfur protein precursor, hydrogenase, monoheme cytochrome subunit, quinol:cytochrome C oxidoreductase subunit II, and quinol:cytochrome c oxidoreductase membrane protein. Transcription levels of these genes were 4.1∼8.6-fold higher in cells in iron-rich compared to iron-limited conditions (Figure 4A). Besides the *bc1* gene cluster, there was a cytochrome *cbb3* gene cluster (Figure 4B) consisting of 8 genes encoding oxidase subunit I/II, subunit III, copper exporting ATPase, copper tolerance proteins, assembly and maturation. The expression of these genes in the *cbb3* gene operon under iron-rich condition was 1.0∼3.1-fold of that compared to the iron-limited conditions (Figure 4B).

**Table 2.** The selected top up- and down-regulated genes determined by RNA-Seq.

| Locus tag | Fold change<br>(Log<sub>2</sub>) | Gene product description |
| --- | --- | --- |
| **Up-regulation** |  |  |
| EAAG1_05702 | 3.10 | Quinol:cytochrome c oxidoreductase |
| EAAG1_09677 | 3.08 | Opacity associated protein (OapA), hypothetical protein |
| EAAG1_09672 | 3.06 | Opacity associated protein (OapA), hypothetical protein |
| EAAG1_05697 | 3.03 | Quinol:cytochrome c oxidoreductase iron-sulfur |
| EAAG1_18235 | 3.02 | Cytochrome c oxidase subunit |
| EAAG1_17456 | 2.89 | Hypothetical protein |
| EAAG1_18230 | 2.55 | Cytochrome c class protein |
| EAAG1_05677 | 2.54 | Quinol:cytochrome c oxidoreductase |
| EAAG1_11612 | 2.52 | Hypothetical protein |
| EAAG1_03478 | 2.52 | Succinate dehydrogenase (or fumarate reductase) |
| **Down-regulation** |  |  |
| EAAG1_10377 | -6.13 | Monooxygenase |
| EAAG1_10367 | -5.77 | Siderophore synthetase component |
| EAAG1_10382 | -5.69 | Putative L-2,4-diaminobutyrate decarboxylase |
| EAAG1_10372 | -5.53 | Hypothetical protein |
| EAAG1_00155 | -5.34 | Hypothetical protein |
| EAAG1_00740 | -4.76 | Putative outer membrane receptor |
| EAAG1_14461 | -4.65 | Hypothetical protein |
| EAAG1_12352 | -4.58 | Hemin-degrading family protein |
| EAAG1_12337 | -4.58 | Transport system permease |
| EAAG1_00245 | -4.06 | TonB-dependent siderophore receptor |

**Figure 4.**
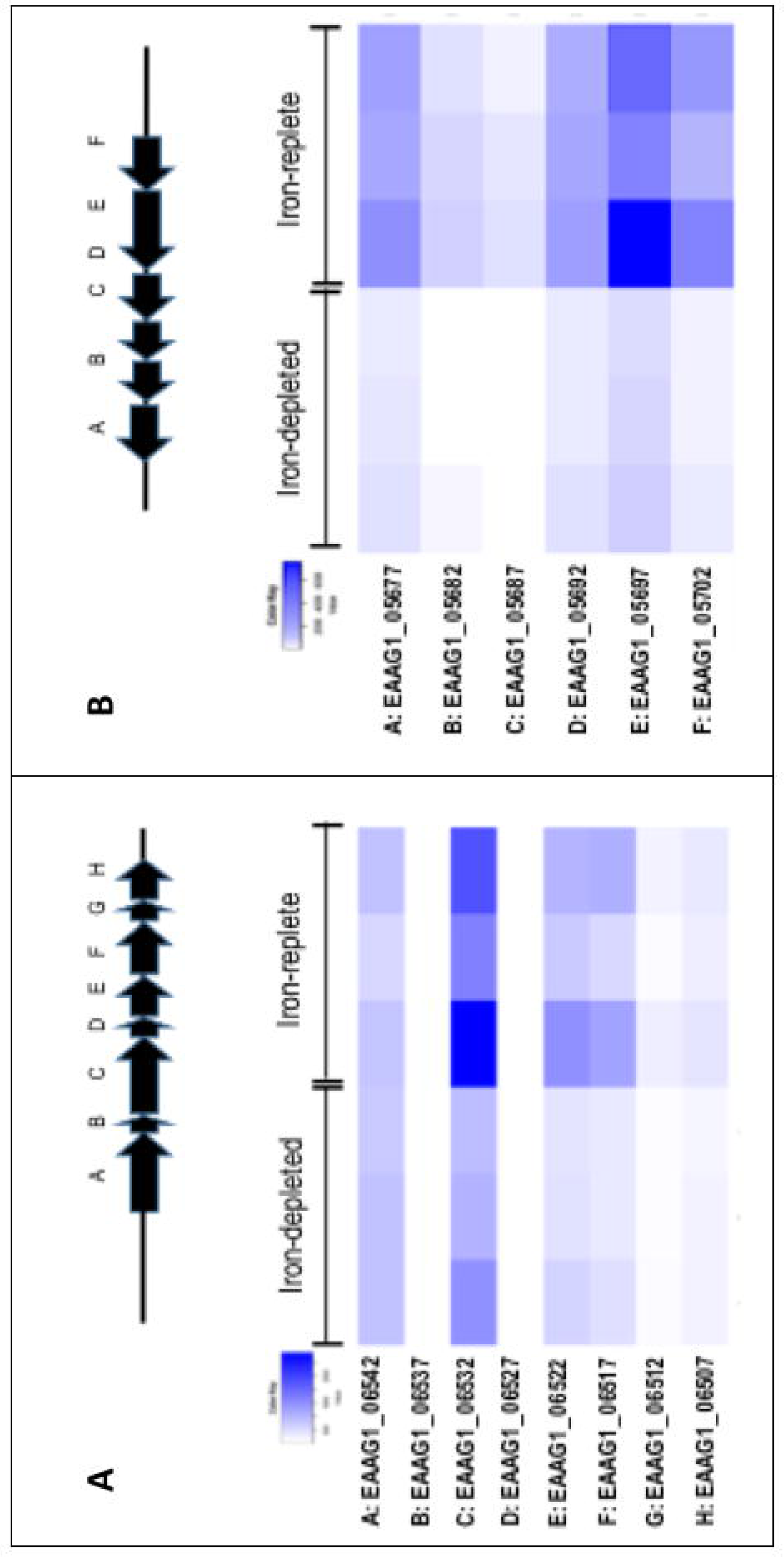
Genes encoding respiratory chain complex *bc1* and *cbb3* in response to iron availability. A) The scheme of genome organization in the *bc1* operon, with heat map of the cytochrome *cbb3* gene expression under the low iron and high iron conditions. (B) The scheme of genome organization in the *bc1* operon, with heat map of the cytochrome *cbb3* gene expression under the low iron and high iron conditions.

### Hemolysin gene expression *in vivo* and *in vitro*

Expression of luciferase in SCH908 (Figure 5A) was 13.9-fold and 20.8-fold higher in iron-low than that in iron-rich cultures at 22°C and 37°C, respectively. Moreover, under iron-low conditions, the relative reporter activity in SCH908 cells cultured at 22°C was 29.1-fold higher than at 37°C; under iron-rich conditions, luciferase activity in SCH908 cells grown at 22°C was 43.2-fold higher than at 37°C. When SCH908 cells were introduced into female mosquitoes by the oral feeding route, luciferase activity measured from dissected guts of sugar-fed mosquitoes was 2.4-fold higher than it was from guts of blood-fed ones, indicating that this promoter was regulated by relative iron availability (Figure 5B). Our results (Figure 5C) further demonstrated that SCH908 cell density in blood-fed *A. stephensi* mosquitoes was 3.3-fold higher than that in the mosquitoes fed with sugar meal (p < 0.05), indicating *E. anophelis* utilized iron from animal blood cells and other nutrients for fast growth in insect host.

**Figure 5.**
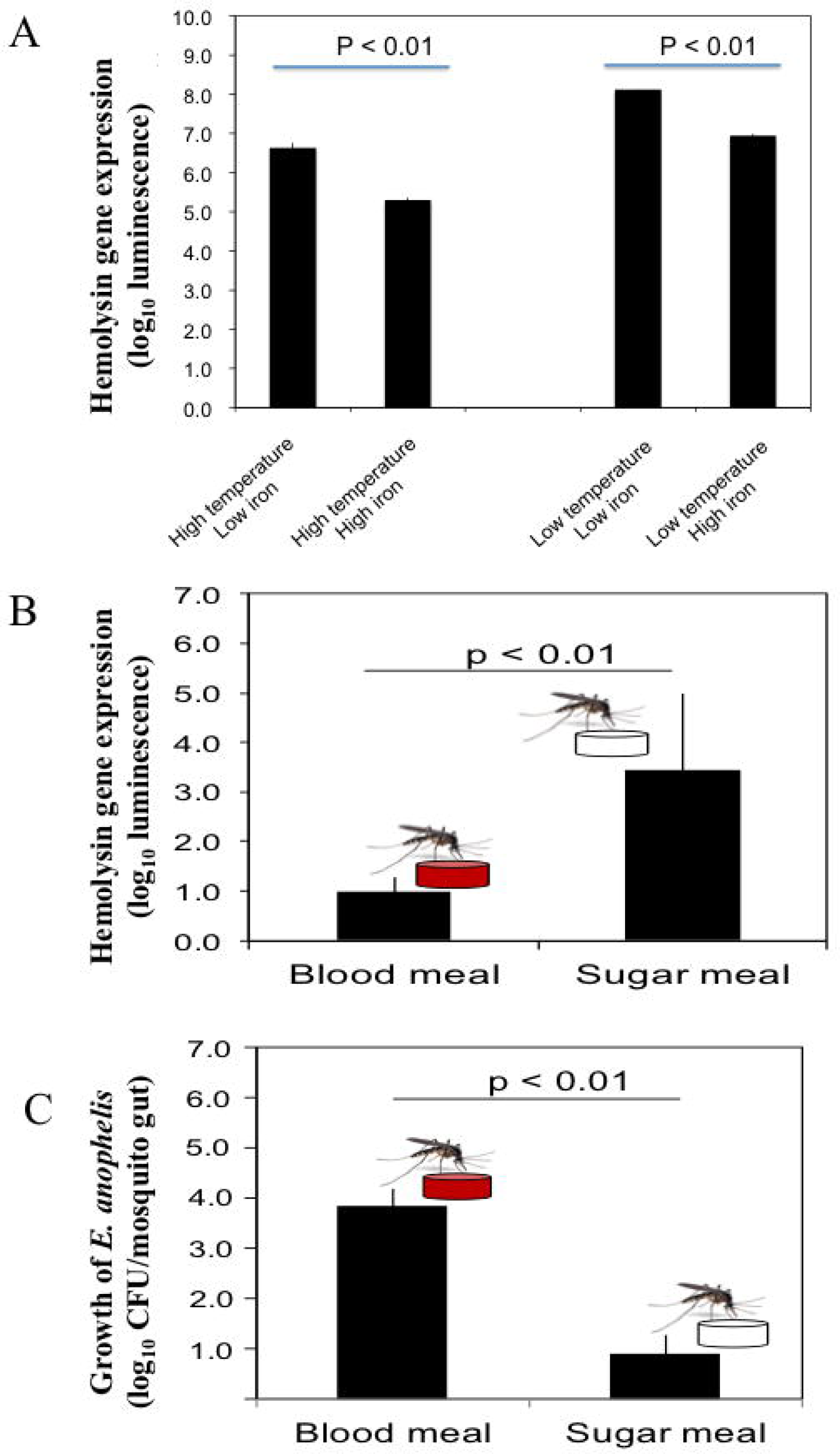
Response of hemolysin genes to temperature, iron stress and diet changes in mosquitoes. (A) Effects of iron and temperature on hemolysin gene expression by *E. anophelis in vitro*. The high or low temperature was 37°C or 22°C, respectively. High-iron medium was established with 12 µM of iron (final concentration), while the low iron medium was established with 200 mM of 2,2-dipyridyl (final concentration). Values are means ± standard deviation. (B) Comparison of hemolysin gene expression in *E. anophelis* when mosquitoes were fed sugar and blood meals. *A. stephensi* were fed with 10% sucrose supplemented with *E. anophelis* for 24 hours (NanoLuc reporter strain). For blood meals, mosquitoes were fed bovine blood through a membrane. Values are means ± standard deviation. (C) Density of *E. anophelis* in mosquitoes given a blood meal or sugar meal. Cell growth was expressed relative to that measured with sugar meal as 100%. Values are means ± standard deviation. Differences were significant at *P*<0.05.

### Hemolysis of erythrocytes and egg production

*E. anophelis* produced alpha-hemolysin on blood agar (Figure S3A). Visual inspection of liquid culture with or without *E. anophelis* Ag1 cells showed less heme color in the former condition, indicating heme utilization by cells (Figure S3B). Under conditions with or without *E. anophelis* Ag1 cells, 14% and 34% of the initial erythrocytes (day 0) were disrupted after 2- and 4-day incubation *in vitro*, respectively, while in cell-free controls erythrocyte density decreased by 5% and 15% of initial RBC count, respectively (Figure 6A). In female mosquitoes not treated with erythromycin or treated with it but after having been fed *E. anophelis* Ag1 via a sugar meal, fecundity averaged 30 eggs per *A. stephensi* (Figure 6B). By contrast, there was an average of 60 eggs per *A. stephensi* after having been fed *E. anophelis* Ag1 in a sugar meal, but not given erythromycin, indicating that *E. anophelis* Ag1 contributed to host fecundity (Figure 6B).

**Figure 6.**
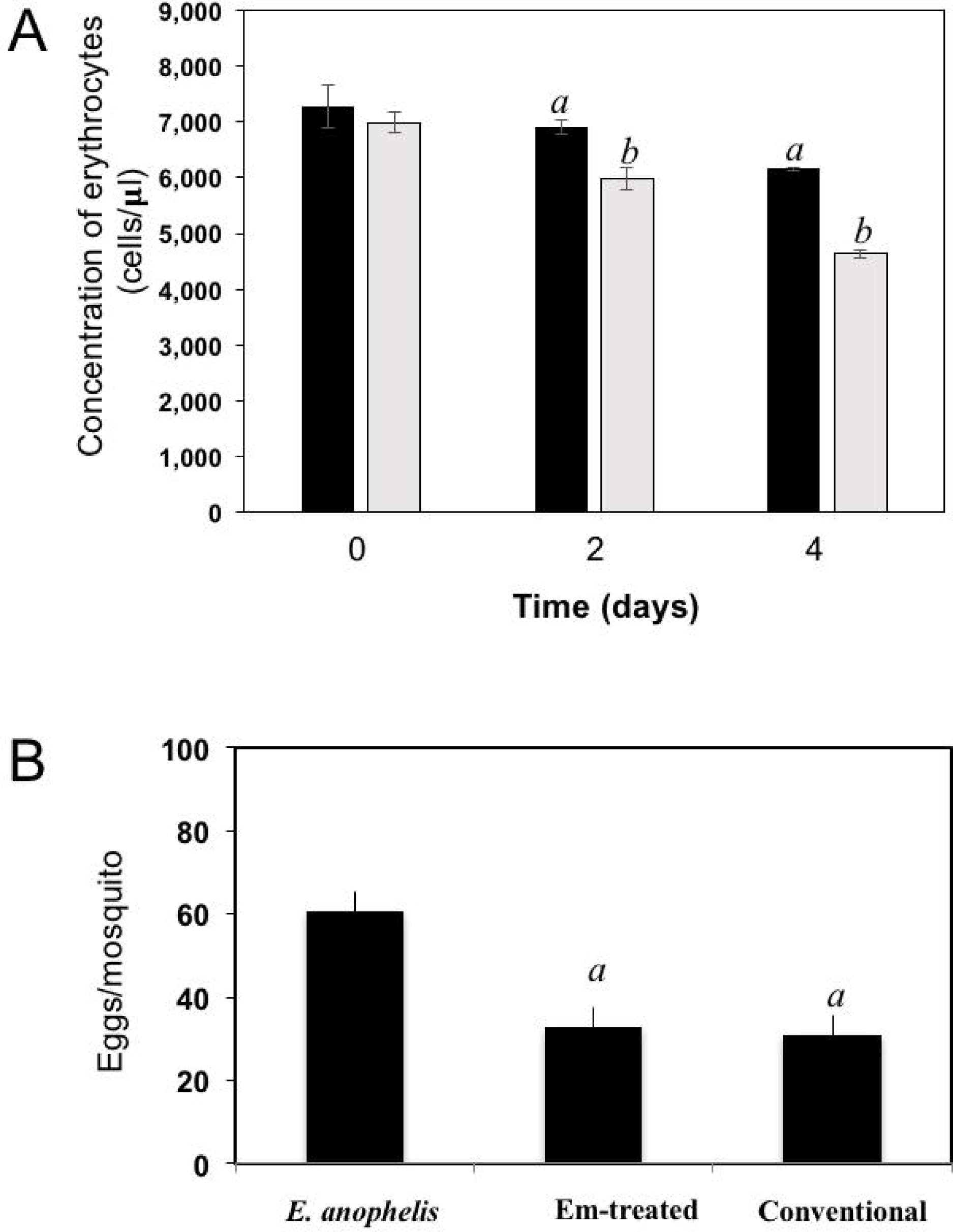
Effects of added *E. anophelis* cells on red blood cell lysis and mosquito fecundity. (A) Time course of the digestion of red blood cells by *E. anophelis*. Values are means ± standard deviation. Comparisons that were significantly different (*P* < 0.05) were concentrations of red blood cells incubated without *E. anophelis* addition on day 2 and day 4 versus day 0 (indicated by “*a*”) or incubated with *E. anophelis* on day 2 and day 4 versus day 0 (indicated by “*b*”), respectively. (B) Fecundity of *A. stephensi* after a bovine blood meal given by membrane feeding. Comparisons that were significantly different (*P* < 0.05) were the eggs produced by mosquitoes previously provided *E. anophelis* in a sugar meal, *E. anophelis* in sugar meal supplemented with erythromycin versus sugar only (indicated by “*a*”), respectively.

### Mutagenesis analysis of the siderophore synthesis genes and biofilm formation

Aerobactin siderophore biosynthesis genes including *iucA/iucC, iucB* and *iucD* were significantly upregulated under iron-poor conditions; their expression level was respectively 53.6, 45.2 and 69.2-fold higher than that in cells held in iron-rich conditions (Table S3). When the three genes Δ*iucA_iucC/iucB/iucD* were intact in wild type *E. anophelis* Ag1, the expected gene fragments were amplified in PCR with appropriate primers; when deleted, these gene fragments were absent, confirming successful construction of deletion mutants (Figure 7A, B). No significant siderophore production in WT was observed in cells grown in iron-rich conditions (Figure S4). However, the blue CAS solution turned to an orange/brown color and the OD_630nm_ decreased when exposed to filtered supernatants of *E. anophelis* grown in ABTGC media with no iron addition, demonstrating significant siderophore activity (Figure S4). *E. anophelis* produced approximately 27 µM of siderophore when deferoxamine was used as an iron siderophore standard (Figure S4). The supernatants of SCH1065 (Δ*iucA_iucC/iucB/iucD*) from the iron-replete culture had a similar OD_630nm_ to the control (no inoculation) when mixed with CAS solution (Figure S4), showing that the function of siderophore synthesis was impaired. The growth of SCH1065 (revealed by OD_600nm_) was comparable to the WT in the first 4 hours under iron-replete conditions (Figure 7C), indicating that *E. anophelis* recycled intracellular iron or scavenged iron from the media using other iron uptake pathways (e.g., direct uptake of available Fe^2+^). However, growth stopped after 4-hour incubation in SCH1065 cells when the culture medium was iron-poor. Under the same conditions, the WT cell density continued to increase and was significantly higher than the mutant cells after 8 h. When cultured under iron-rich conditions, both mutant and WT cells grew to stationary phase at 8 h. However, the final cell density in the mutant was slightly lower than that in WT (Figure 7C).

**Figure 7.**
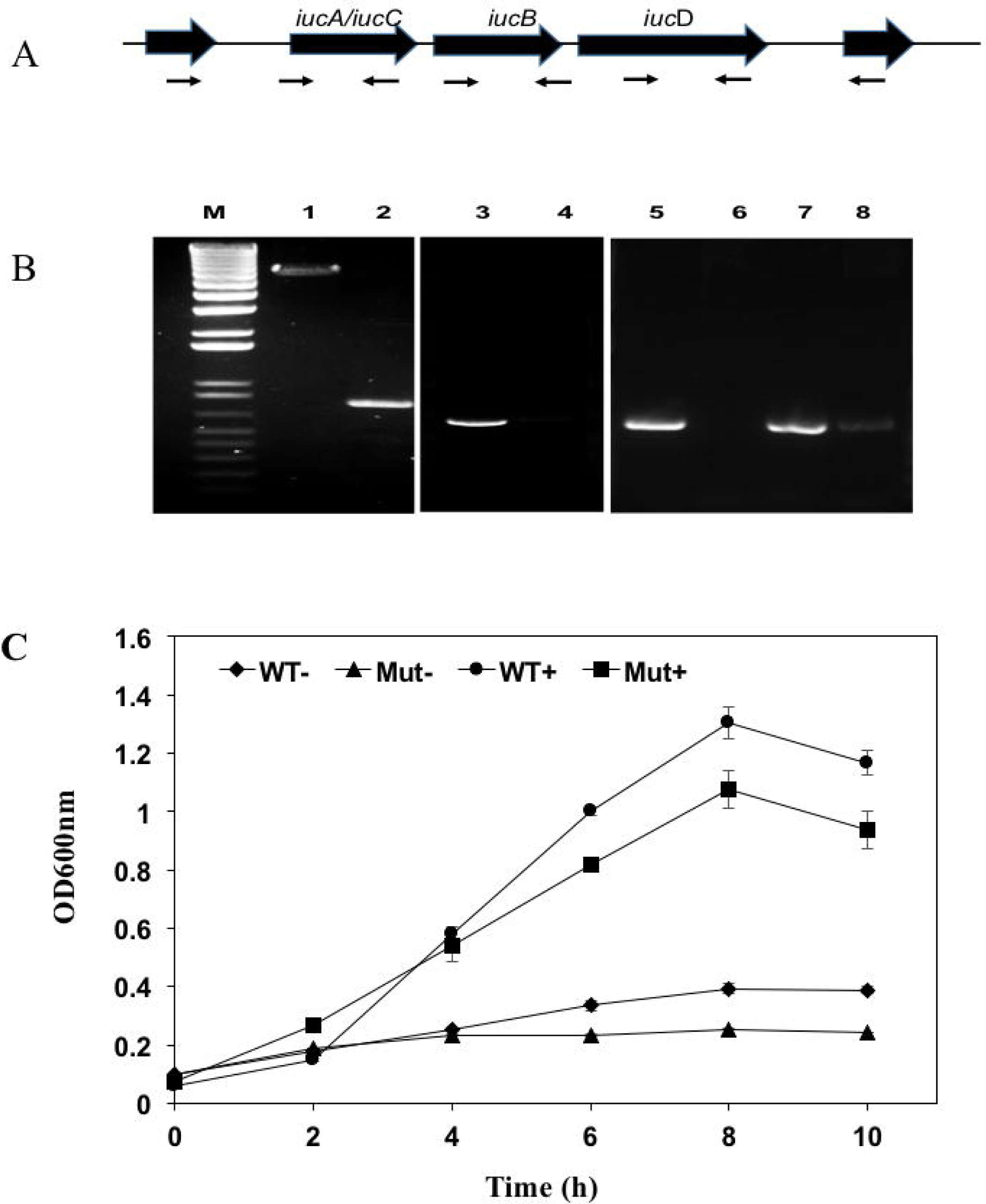
Deletion of the siderophore synthesis gene cluster (Δ*iucA_iucC/iucB/iucD*) led to impaired growth under iron stress conditions. (A) Organization of the *iucA/iucC, iucB* and *iucD* gene cluster in *E. anophelis*. (B) Detection of *iucA/iucC, iucB* and *iucD* genes in the WT and the mutant. Lane M, molecular marker; lane 1, gene fragment of the regions spanning the upstream and downstream of the *iucA/iucC, iucB* and *iucD* gene cluster with the WT genomic DNA by the primers walker295 and walker296; lane 2, gene fragment of the regions spanning the upstream and downstream of the *iucA/iucC, iucB* and *iucD* gene cluster in the mutant by the primers walker295 and walker296; lane 3, gene fragment of *iucA/iucC* amplified with WT genomic DNA by walker297 and walker 298; lane 4, gene fragment of *iucA/iucC* amplified with mutant genomic DNA by walker297 and walker 298; lane 5, gene fragment of *iucB* amplified with WT genomic DNA by walker299 and walker300; lane 6, gene fragment of *iucB* amplified with mutant genomic DNA by walker299 and walker300. lane 7, gene fragment of *iucC* amplified with WT genomic DNA by walker301 and walker302; lane 8, gene fragment of *iucC* amplified with mutant genomic DNA by walker301 and walker302. (C) The growth curve of WT and siderophore synthesis mutant in high-iron and low iron media. WT-, wild-type grown in the low iron media; Mut-, mutant grown in the low iron media; WT+, wild-type grown in the high-iron media; Mut+, mutant grown in the high-iron media.

When wild-type cells were cultured in LB broth, only 30% of the original cells were viably retained after the bacteria were treated with 20 µM of H_2_O_2_ for 20 min (Figure 8A). By contrast, 60% of viable cells were recovered in *Elizabethkingia* pre-grown in LB media with the addition of FeCl_3_ or hemin (Figure 8A), showing that the preloaded cellular iron is critical for bacterial tolerance of the H_2_O_2_ toxicity. Cells grown in hemoglobin had an even better protection from H_2_O_2_ toxicity with a bacterial surviving rate of up to 80% (Figure 8A). Nearly all (97.5%) of cells grown under iron-poor conditions died, indicating that iron-depleted cells were highly susceptible to H_2_O_2_ toxicity (Figure 8A). Further, only 1.1% of the original cells were retained when the siderophore mutant was grown in LB media (Figure 8B). We were only able to recover 0.037% of the original cells when the mutant was cultured in iron-poor medium, showing that cells without efficient uptake of iron were more susceptible to H_2_O_2_ than WT (Figure 8B). Incubation of mutants with hemin, Fe^3+^ or hemoglobin retained at least 0.2, 4% and 3.3% of the original cells without H_2_O_2_ treatment, indicating that iron is important for protection from H_2_O_2_ toxicity (Figure 8B).

**Figure 8.**
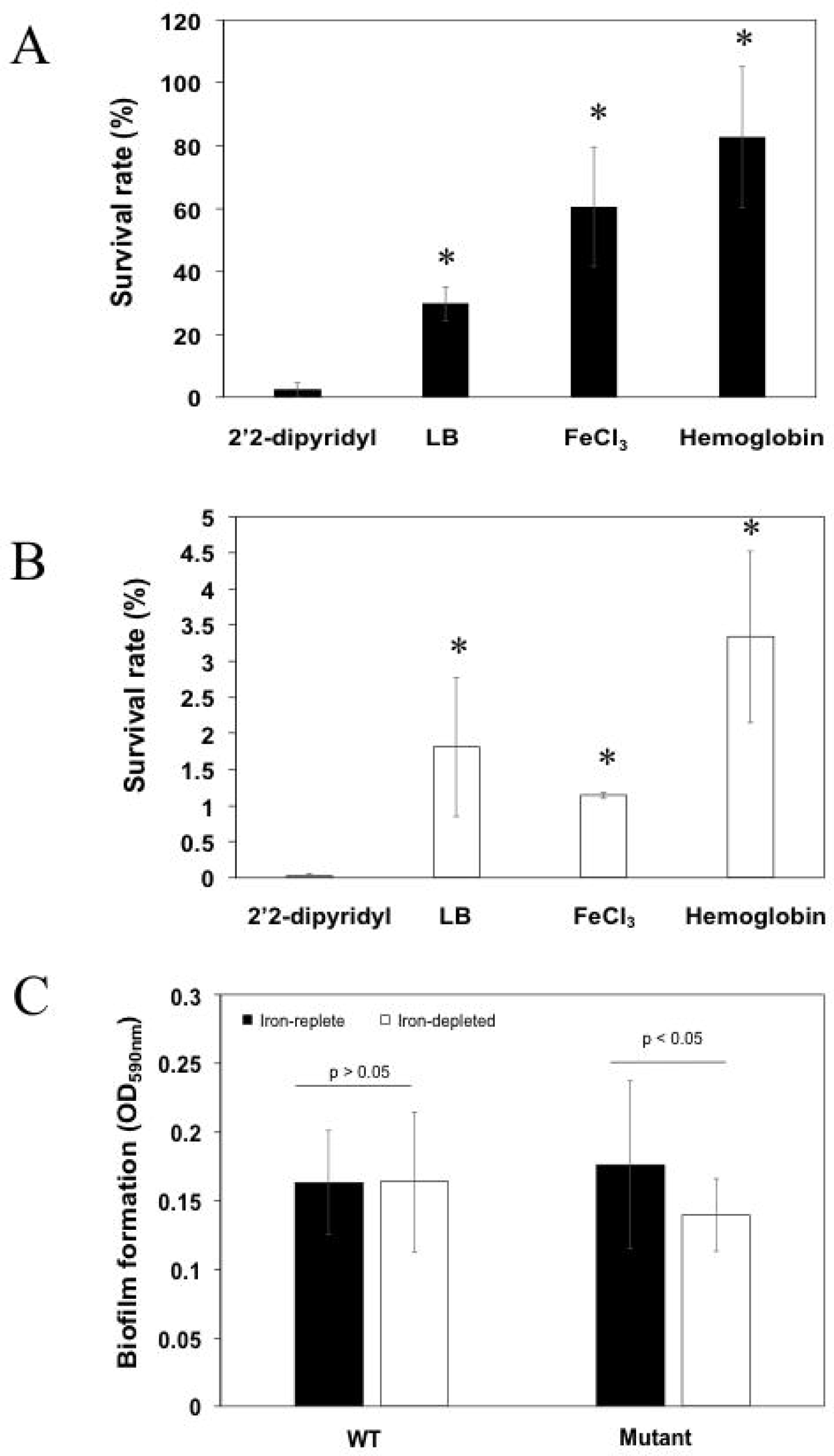
Deficiency in siderophore synthesis was susceptible to H_2_O_2_ damage and led to the attenuated biofilm formation. (A) The survival rate of WT cells in low iron media (LB added with 2’2-dipyridyl), LB media, high iron media, and LB media supplemented with hemoglobin. Values are means ± standard deviation. Asterisks indicated there was a significant difference (*P* < 0.05). (B) The survival rate of mutants in low iron media (LB added with 2’2-dipyridyl), LB media, high iron media, and LB media supplemented with hemoglobin. Values are means ± standard deviation. Asterisks indicate there was a significant difference (*P* < 0.05). (C) Biofilm formation of the WT and mutant cells grown in the low iron and high iron media. Values are means ± standard deviation.

Biofilm formation in the mutant SCH1065 was only 71.8% of the WT when grown in iron-depleted media (Figure 8C). However, the deficiency in the siderophore synthesis (Figure 8C) did not affect biofilm formation if the cultures (WT and mutant) were grown in iron-replete media (p > 0.05).

## Discussion

The female mosquito gut is characterized by slightly acidic pH with a tendency to alkalinity after blood meals, a small volume, and frequent nutritional fluctuations (41). Iron availability contributes importantly to the assemblage of microorganisms in the mosquito midgut (3, 19, 42). Bacterial inhabitants of the adult mosquito midgut experience extreme conditions with relationship to iron availability, either having insufficient supply prior to and between blood meals, or excessive free iron and associated free radicals during and after blood meals (42, 43). Our results showed that *E. anophelis*, a commensal organism of the mosquito midgut, responds to these extreme conditions and utilizes multiple iron uptake pathways, TBDTs, and iron chelating proteins in response to iron stress. We further demonstrated that siderophores are crucial for iron uptake, alleviation of oxidative damage by H_2_O_2_, and biofilm formation. *E. anophelis* facilitated RBC lysis with several hemolysins, which possibly contributed to the mosquito’s fecundity. Collectively, our results provide new insights into the colonization and survival mechanisms for this predominant commensal bacterium in the mosquito gut.

A global gene response occurred in response to iron stress in *E. anophelis* and involved least 9% of the total transcribed genes. In particular, the expression level of several genes involved in “the respiratory chain complex” was notably affected by iron availability. Bacteria utilize several iron-storage proteins (DPS, ferritins and/or bacterioferritin) to remove excessive iron that produces radicals and the oxidative damage inside the cells (22, 44, 45). However, *de novo* synthesis of iron storage proteins is slow and energy-consuming to manage a sudden, high concentration of free iron (22). The expression of iron storage protein genes is frequently detected during stationary growth phase (regulated by transcription sigma factor σ^54^) (45, 46). Incorporating excessive free iron into proteins/enzymes may be a better strategy than *de novo* synthesis of iron storage proteins, when a sudden high load of iron is encountered, because it allows bacteria to remove reactive oxygen radicals quickly, minimizing damage to the cell (47-49). Among the iron-containing proteins (using iron as cofactors), the ETC components are good candidates because they utilize iron/heme as the cofactors and are highly demanded to maintain the cell metabolism (50). Besides the ETC components, expression of genes involved in TCA cycle and/or Fe-S protein synthesis was observed to be significantly elevated in the high iron cells. Guo et al. (2017) also demonstrated that the transcription level of ETC genes was significantly higher (up to 5-fold) in iron-rich cells than that in iron-poor cells in *R. anatipestifer* CH-1 (51). The transcription level of ETCs (such as cytochrome-c peroxidase, cytochrome c oxidase subunit CcoP and cytochrome c oxidase subunit CcoN) was remarkably lower in H_2_O_2_-treated cells than in untreated *E. anophelis* (38). *E. anophelis* dramatically decreased iron uptake, likely to avoid more intracellular ROS production when high oxidative stress (presented here experimentally by H_2_O_2_ treatment) is encountered in the environment (38). Therefore, the rapid conversion of toxic free iron into non-toxic forms, and minimizing iron uptake, substantially contributes to cellular tolerance to the high oxidative stress.

Hemolysin production has been widely reported in many pathogens (52-55). *E. anophelis* produced *α*-hemolysin(s) to facilitate digestion of animal erythrocytes *in vitro*. Restriction of environmental iron increased hemolysin gene transcription, demonstrating that they were actively involved in iron metabolism. Once erythrocytes are lysed, iron/heme becomes available in the mosquito midgut (56). In our experiments, *E. anophelis* dramatically lowered hemolysin gene expression, perhaps to avoid excessive hemoglobin release. By contrast, in *S. marcescens*, expression of hemolysin genes greatly increased when iron in the medium was limited (57). In *Yersinia ruckeri*, the promoter activity of these genes was regulated by both iron concentration and temperature (58). Infection was more efficient at low temperature (18°C), which induced higher expression of hemolysin, protease (*Yrp1*), ruckerbactin and other toxin genes than at elevated temperature (37°C) (58). Our results demonstrate that expression of hemolysin genes was remarkably depressed by increasing environmental temperature though the bacteria grow faster at 37°C. For *Elizabethkingia*, the temperature-dependent modulation of hemolysin in the blood feeding course may be similar to the scenario encountered by the *Y. ruckeri* in the blood stream (58). When the female mosquito ingests blood cells from warm-blooded animals, the midgut epithelial cells respond with rapid expression of heat shock proteins (59). The higher temperature could be a signal for *Elizabethkingia* in the midgut to decrease hemolysin synthesis. However, further experiments are warranted to test this hypothesis.

Supplementation of *E. anophelis* in the diet of adult female *A. stephensi* increased fecundity. Gaio et al. (2011) investigated the effects of the gut bacteria on the blood digestion and egg production in *A. aegypti* (56). Elimination of the gut bacteria slowed digestion of erythrocyte protein components (56). It is well established that blood components induce release of insulin-like peptides (ILPs) and ovary ecdysteroidogenic hormone (OEH) in mosquitoes and stimulate ecdysone production (60-62). Mosquitoes require ecdysone and ILPs to produce yolk proteins which are incorporated into primary oocytes forming mature eggs (60, 61). Consequently, retarded release of essential nutrients involved in the vitellogenic cycle due to the removal of gut microbiota possibly affected oocyte maturation, resulting in the production of less viable eggs (60, 61). To efficiently digest proteins from whole blood cells, the anautogenous mosquito hosts (such as *A. aegypti* and *A. stephensi*) and their associated gut bacteria are required to work in synergism (62). Remarkably, there are redundant peptidases/proteinases in *E. anophelis* (at least 77, data not shown). Some peptidase genes were differentially up-regulated by the iron (data not shown), possibly contributing to blood protein digestion and facilitating the increased fecundity observed here.

When mosquitoes are only fed a sugar meal, *Elizabethkingia* may access ferrous iron through ferrous transporters, ferric iron using siderophores, and other iron sources by exosiderophore (i.e. ferrichrome, see below) in the gut. Due to the aerobic environment in mosquito gut, ferric iron may be one of the prevalent iron species in sugar-fed mosquito. However, the bioavailability of ferric iron in water is extremely low (about 10^-17^ M) (63). Secretion of efficient siderophores such as aerobactin from the mosquito-associated bacteria is important for competing for the very limited iron resource in the mosquito midgut (38). The siderophore synthesis deficient mutants were particularly vulnerable to oxidative damage, indicating that siderophore production in *E. anophelis* can protect the cells from H_2_O_2_ damage (38). The ability to remove free radicals is seriously compromised if hosts cannot produce iron-containing peroxidases and catalases. As an opportunistic and emerging pathogen, *Elizabethkingia* may utilize aerobactin as an important virulence factor. Among the different siderophores (i.e., aerobactin, salmochelin, enterobactin and yersiniabactin), aerobactin was one of the most important virulence factors in systemic infections (64).

Hemin, Fe^3+^ or hemoglobin (Hgb) attenuated cytotoxicity by H_2_O_2_ (38). Addition of hemoglobin to *E. anophelis* NUHP1 significantly enhanced H_2_O_2_ resistance (38). Exposure to H_2_O_2_ accumulates reactive oxygen species (ROS) and nitric oxide (NO), decreases mitochondrial membrane potential, and triggers caspase-3/7 activity in vertebrates (65). Hemoglobin remarkably attenuated the overproduction of ROS and NO, reverted mitochondrial membrane potential, and repressed caspase-3/7 (65). It is possible that the hemin/iron from degraded hemoglobin can immediately be transported into cells and incorporated to some of the heme-containing antioxidative enzymes (such as heme-containing catalases) when bacteria suffer the crisis of strong reactive oxygen radicals (42, 66). Similarly, some of the non-heme proteins (with iron as the cofactor) can remove the ROS (66).

Iron availability impacted biofilm formation early in its development (67). In this study, the siderophore-deficient mutant formed less biofilm than the WT under iron-depleted conditions. Iron-responsive genes such as siderophore synthesis and iron uptake genes were strongly induced by biofilm formation rather than by planktonic growth in *Mycobacterium smegmatis* (68). Further, deficiency in the exochelin biosynthesis or uptake systems led to poor biofilm formation, the viability and cultivability of biofilm cells under iron-limiting conditions (67, 68). Biofilm formation *in vitro* often led to weaker ability to attach to animal cells in *Elizabethkingia*. Thus, our observations here indicate that siderophore synthesis is also important for successful colonization in mosquitoes.

Environmental stress (e.g. pH and temperature), immune defense, as well as nutritional variations for commensal *Elizabethkingia* in mosquitoes with sudden blood meals are similar to those invading the bloodstream (19, 69). Moreover, genome contexts and gene sequences between the mosquito-associated *E. anophelis* and clinical isolates are conserved (9, 12, 30, 38). Thus, the investigation of the molecular mechanisms of the iron metabolism in mosquito-associated *E. anophelis* may also contribute to the understanding of the pathogenesis progress in the same species and the similar organisms.

## Author contributions

SC and EDW conceived the study and participated in its design and coordination. SC and BNN performed the experiments. SC, BKJ and YT conducted the transcriptomics analysis. SC and EDW wrote the manuscript. All authors have read and approved the manuscript.

## Acknowledgments

This project was funded by NIH grant R37AI21884. Authors thank Dr. Mark McBride at University of Wisconsin-Milwaukee and Dr. Jiannong Xu at the New Mexico State University for the plasmid pYT313 and the strain *E. anophelis* Ag1.

