## Supplemental Figures for "*Elizabethkingia anophelis* response to iron stress: physiologic, genomic, and transcriptomic analyses"

Send correspondence to:

Dr. Shicheng Chen

Department of Microbiology and Molecular Genetics

2215 Biomedical and Physical Sciences Building

Michigan State University

567 Wilson Road

East Lansing, Michigan 48824-4320

517-884-5383

***Elizabethkingia anophelis* response to iron stress: physiologic, genomic, and transcriptomic analyses**

Shicheng Chen^1^, Benjamin K. Johnson^1^, Ting Yu^2^, Brooke N. Nelson^1^

and Edward D. Walker^1, 3^

^1^Dept of Microbiology and Molecular Genetics, Michigan State University, East Lansing, MI 48824 USA

^2^Agro-biological Gene Research Center, Guangdong Academy of Agricultural Sciences, Guangzhou, 510640 China

^3^Dept of Entomology, Michigan State University, East Lansing, MI 48824 USA

Running title: transcriptomic analysis of *Elizabethkingia* under iron-stress conditions

**Supplemented Materials**

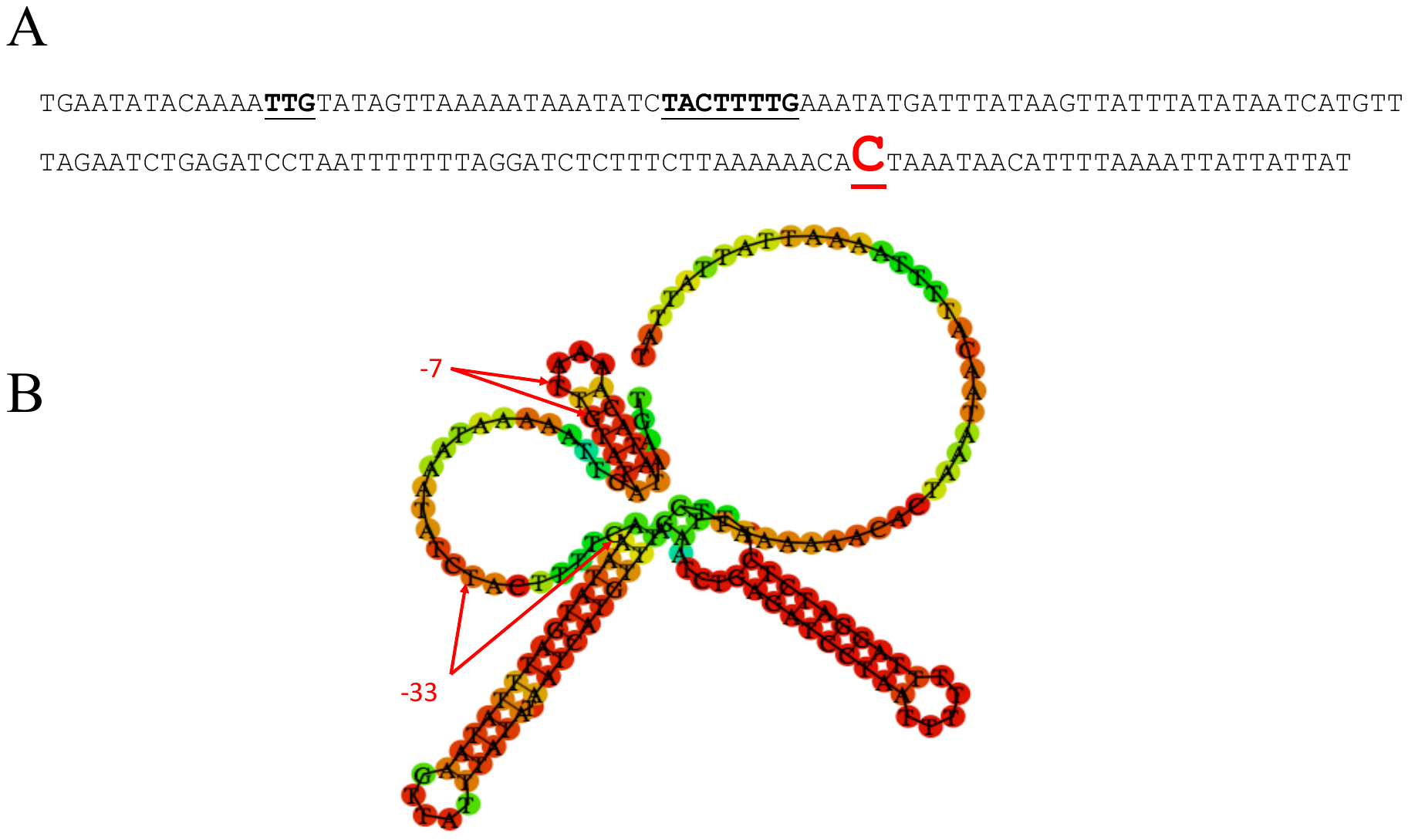

**Figure S1. The transcriptional start site (TSS) for gene of Elilysin2 (EAAG1_11027) determination and promoter prediction.** A) The TSS was determined as C, 29 bp upstream of the translation start site. Red font and underlined. The promoter motifs were highlighted with underline. B) The folding of the promoter regions was predicted with RNAfold (<http://rna.tbi.univie.ac.at)>.

*E. anophelis* grown in LB broth with high or low iron concentration respectively, to an OD600 of about 0.5. Cells were harvested by centrifugation at 4,000 g for 10 minutes at room temperature. Pellets were resuspended in 2 ml of RNAlater (Qiagen, USA) and stored at -80°C. Bacterial cell pellets stored at -80°C in RNAlater (25 ml) were thawed on ice, re-suspended and re-pelleted in 1 ml aliquots for 10 minutes at 4,000 g in a microcentrifuge. The supernatant was removed and 200 μl bacterial lysis buffer (30 mM Tris HCl, pH 8.0, 1 mM EDTA plus 15 mg/ml lysozyme (Sigma, St Louis, MO, USA) and 15 μl proteinase K (20mg/ml; QIAGEN, Valencia, CA, USA) were added to each tube. Samples were incubated at room temperature for 10 minutes, and vortexed for 10 s before and every 2 minutes during the incubation. QIAGEN RLT Plus buffer (750 μl) supplemented with 1 % v/v beta-mercaptoethanol (Sigma) was added to each tube and vortexed briefly to mix. Analysis by 5’ rapid amplification of cDNA ends (5’RACE) was done by using the SMARTer RACE 5’/3’ kit (Clontech, CA) under the manipulation procedures recommended by the supplier. For Pheam, the first cDNA strand was done with a specific primer Walker188 which is based on hemolysin gene sequence (Table 2). The 5’-end region of interest was then amplified by using two sets of primers for regions of heamlysin in a nested PCR. In the first PCR, RT products were used as a template and amplified with primers Walker188 and UPM (Universal Primer Mix). The second PCR (nested PCR) was conducted with primers Walker189 and Nested UPM with the 10-fold dilution of the round 1 PCR product as the template (Table 2). The 5-RACE product with a size of ~450-bp were isolated, purified, ligated into the pGEM-T Easy vector, and sequenced. 5’RACE-PCR showed that the transcriptional start site is C. Immediate 29-bp upstream of the translational start site, there is a conserved promoter motif TTG-N19-TAnnTTTG (ref). The promoter resembles the consensus promoter structure in other *Bacteroidetes*.

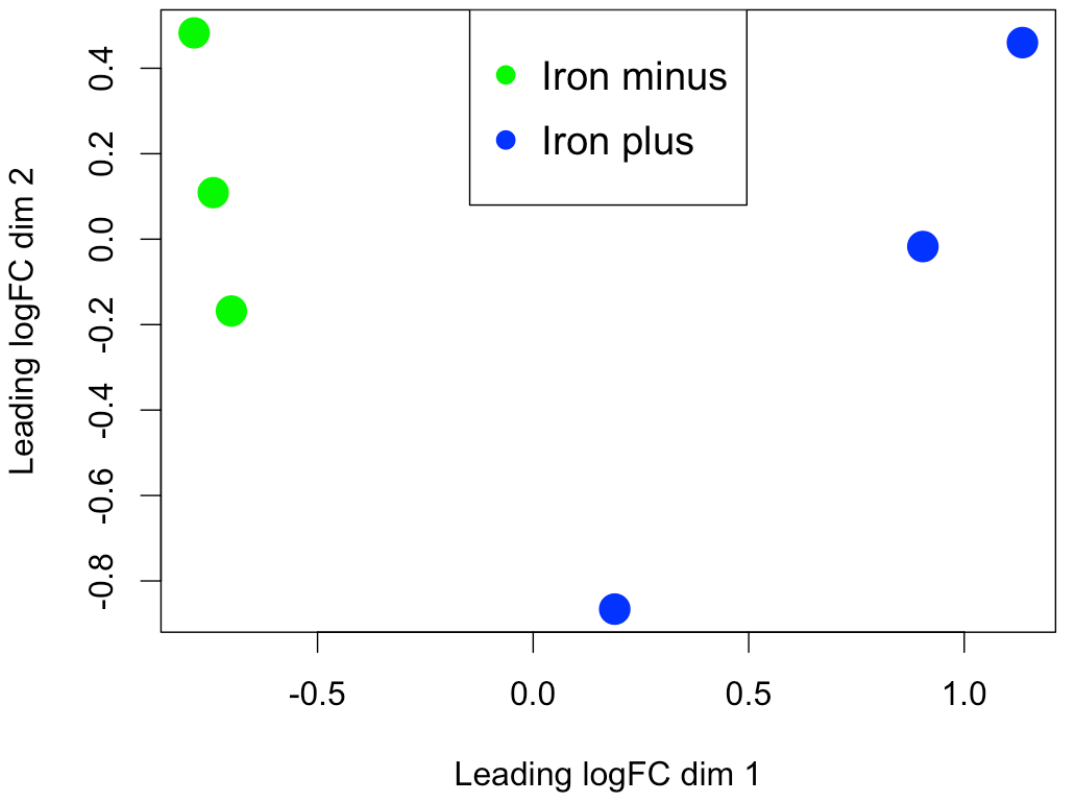

**Figure S2. PCA analysis of the high and low iron RNAseq libraries.** The iron minus indicated low-iron cultures and iron plus indicated the high iron cultures.

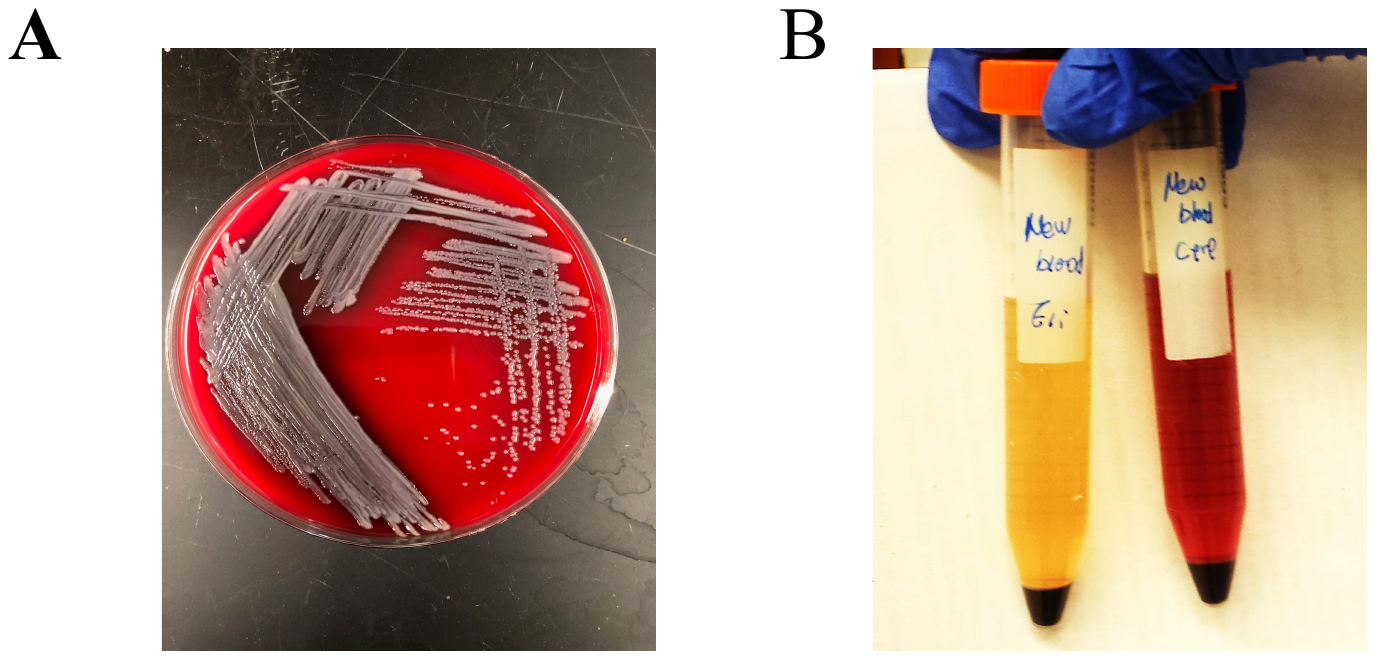

**Figure S3. Demonstration of the alpha-hemolysin production and hemoglobin uptake in *E. anophelis*.** A) The greenish colonies on SBA indicated the alpha-hemolytic activity. B) The left tube was inoculated with *E. anophelis* and the right one is control without inoculation. The vials were rotated for overnight and centrifuged.

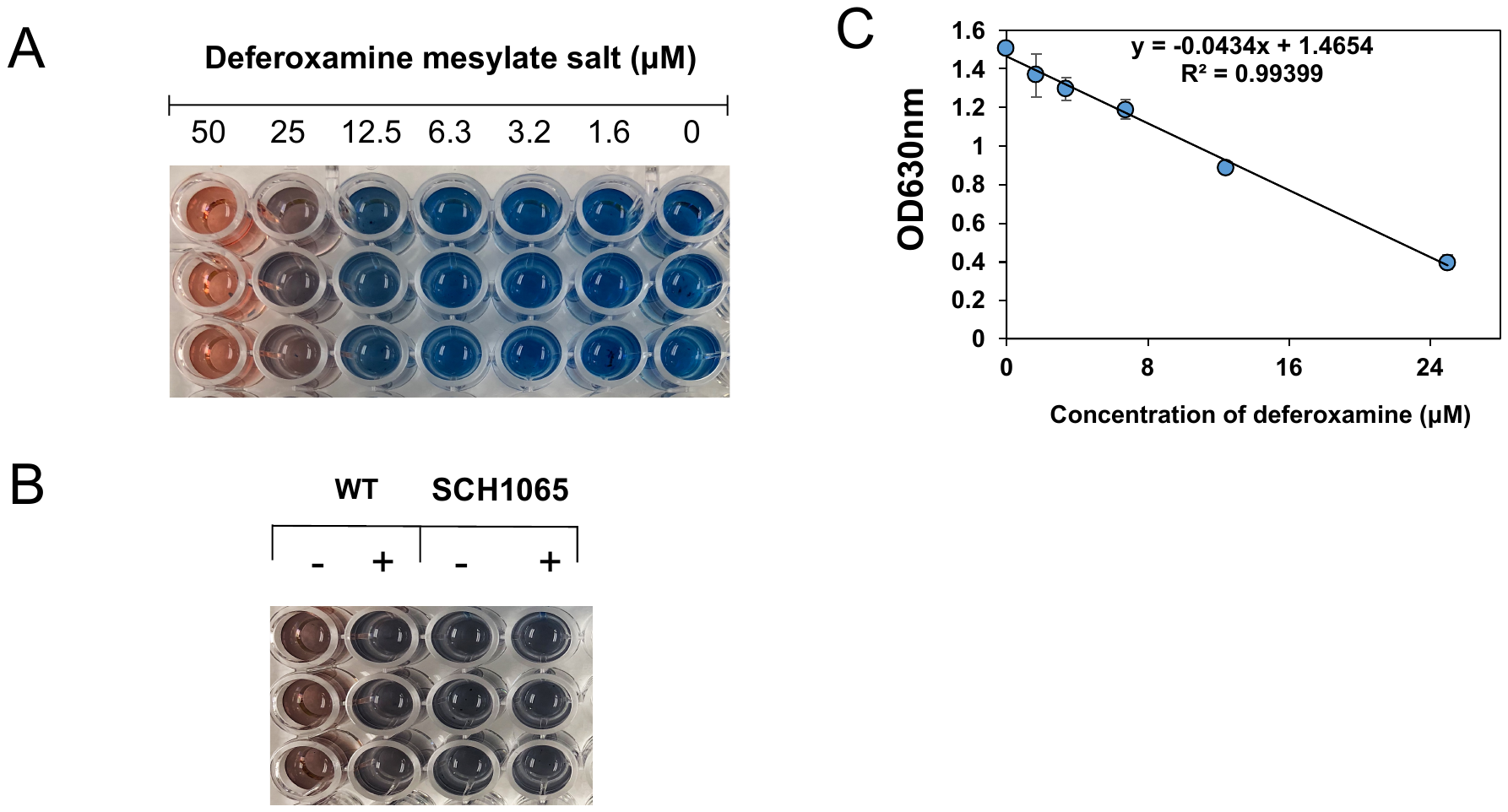

**Figure S4. The siderophore production and determination in WT and mutants of *E. anophelis*.** A) Effects of the adding deferoxamine mesylate on the color change of the iron solution as described in the Methods and Materials. B) The different color change caused by the siderophore production between the WT and the mutant. C) The standard curve for determination of the siderophore production using deferoxamine mesylate.
